## SUPPLEMENTAL_FIGURES for "TARPON - a Telomere Analysis and Research Pipeline Optimized for Nanopore"

**a**

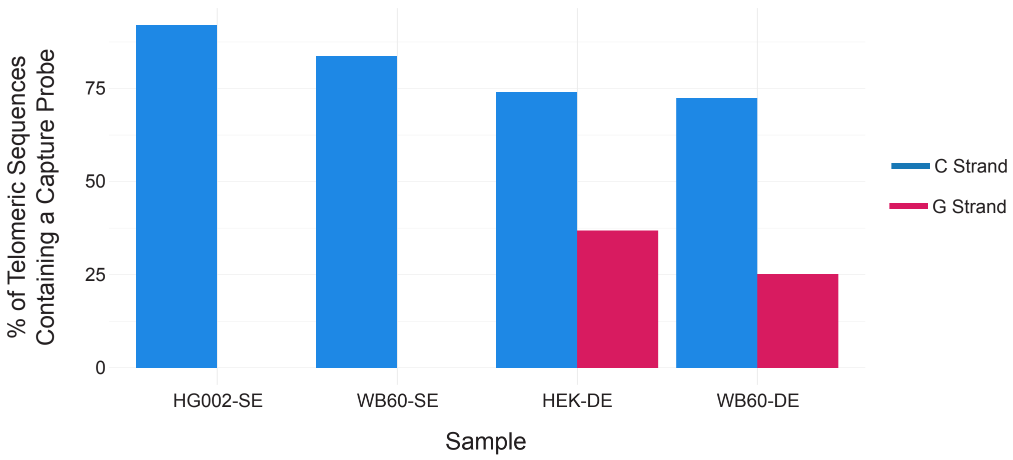


**Deimler, 2025; Supplemental Figure 1**

a

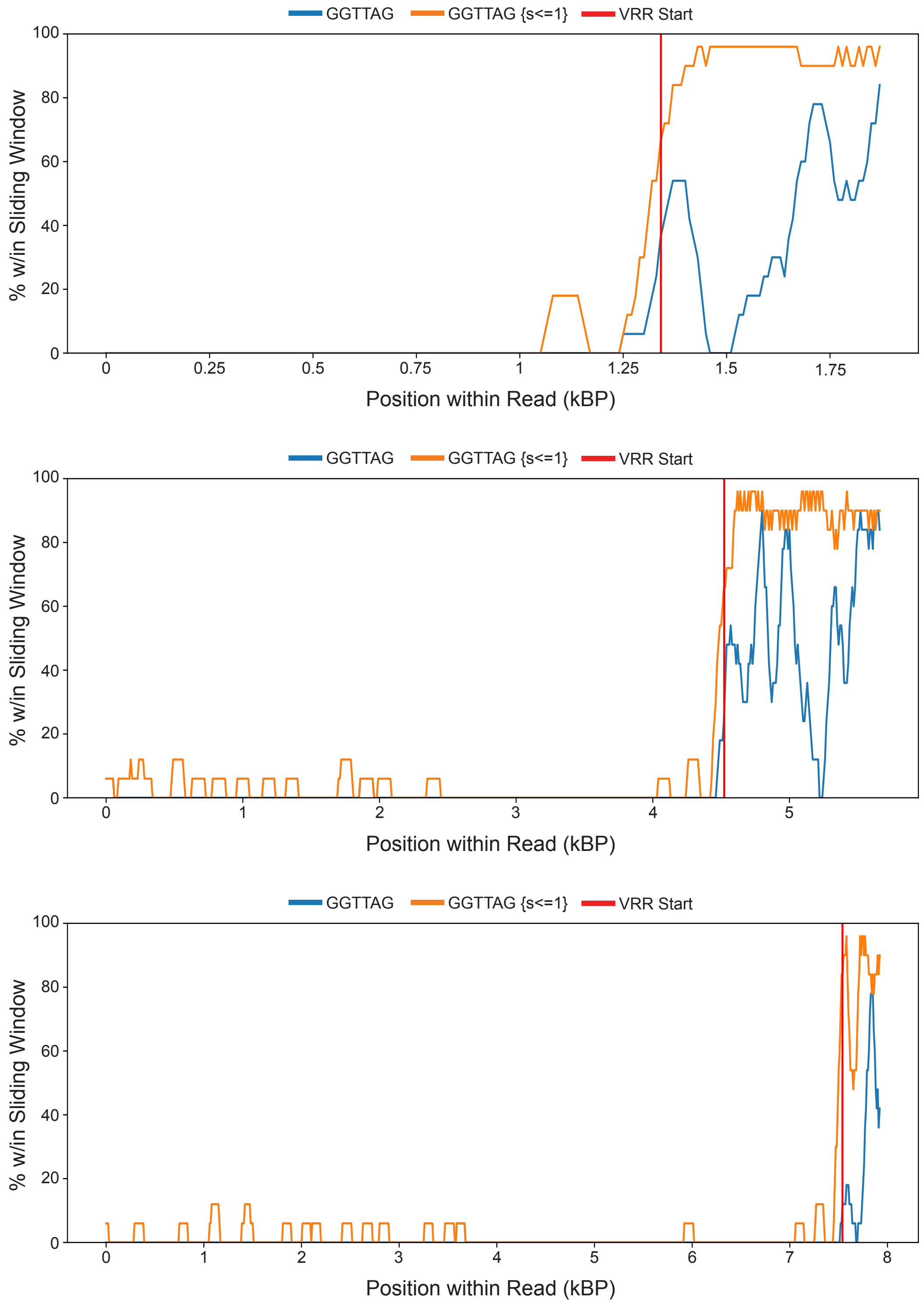


**Deimler, 2025; Supplemental Figure 2**

**a**

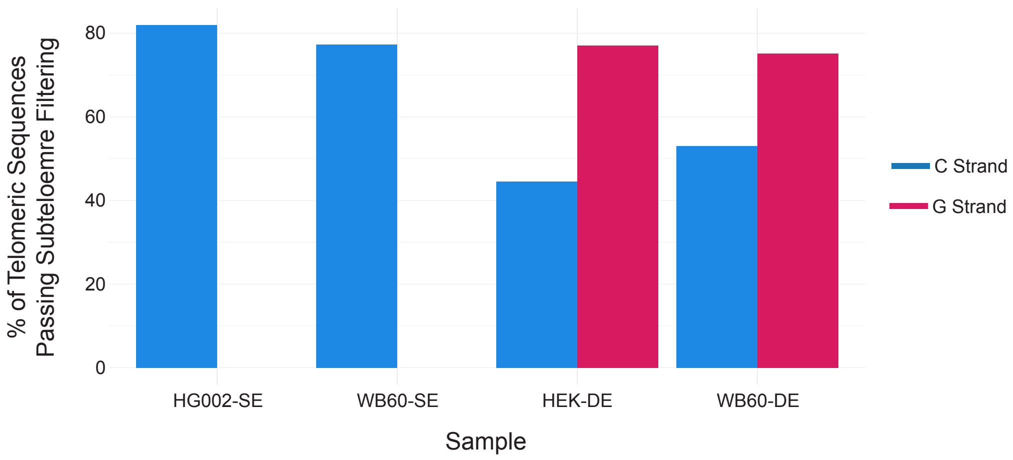


**Deimler, 2025; Supplemental Figure 3**

**a**

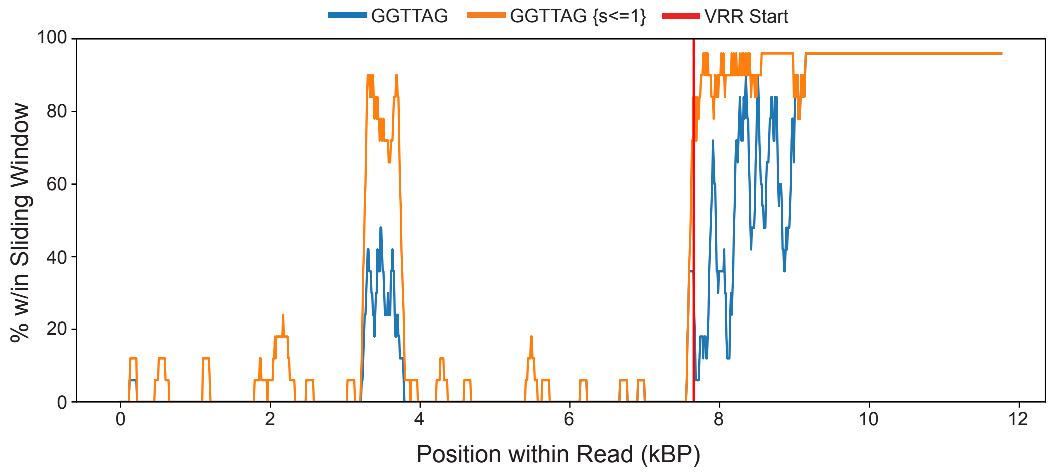
GGTTAG
GGTTAG {s<=1}
VRR Start
% w/in Sliding Window
100
80
60
40
20
0
0
2
4
6
8
10
12
Position within Read (kBP)

**a**

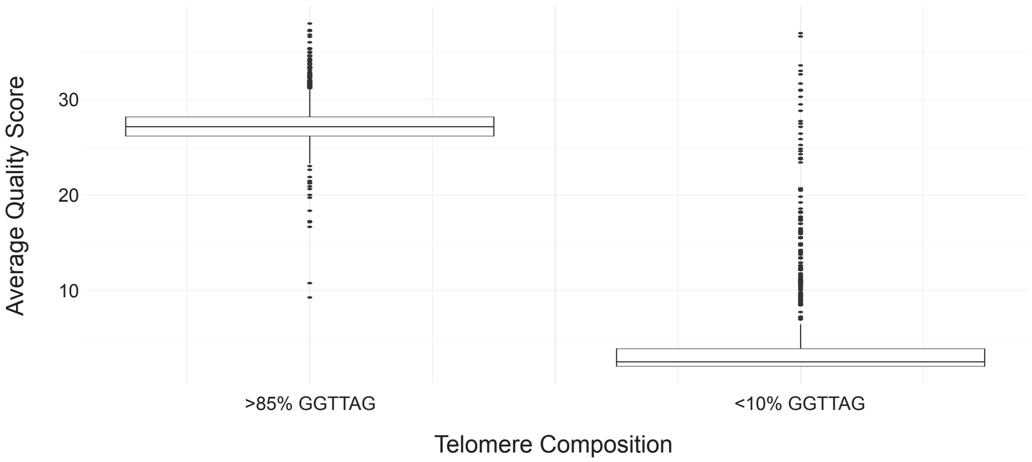


**Deimler, 2025; Supplemental Figure 5**

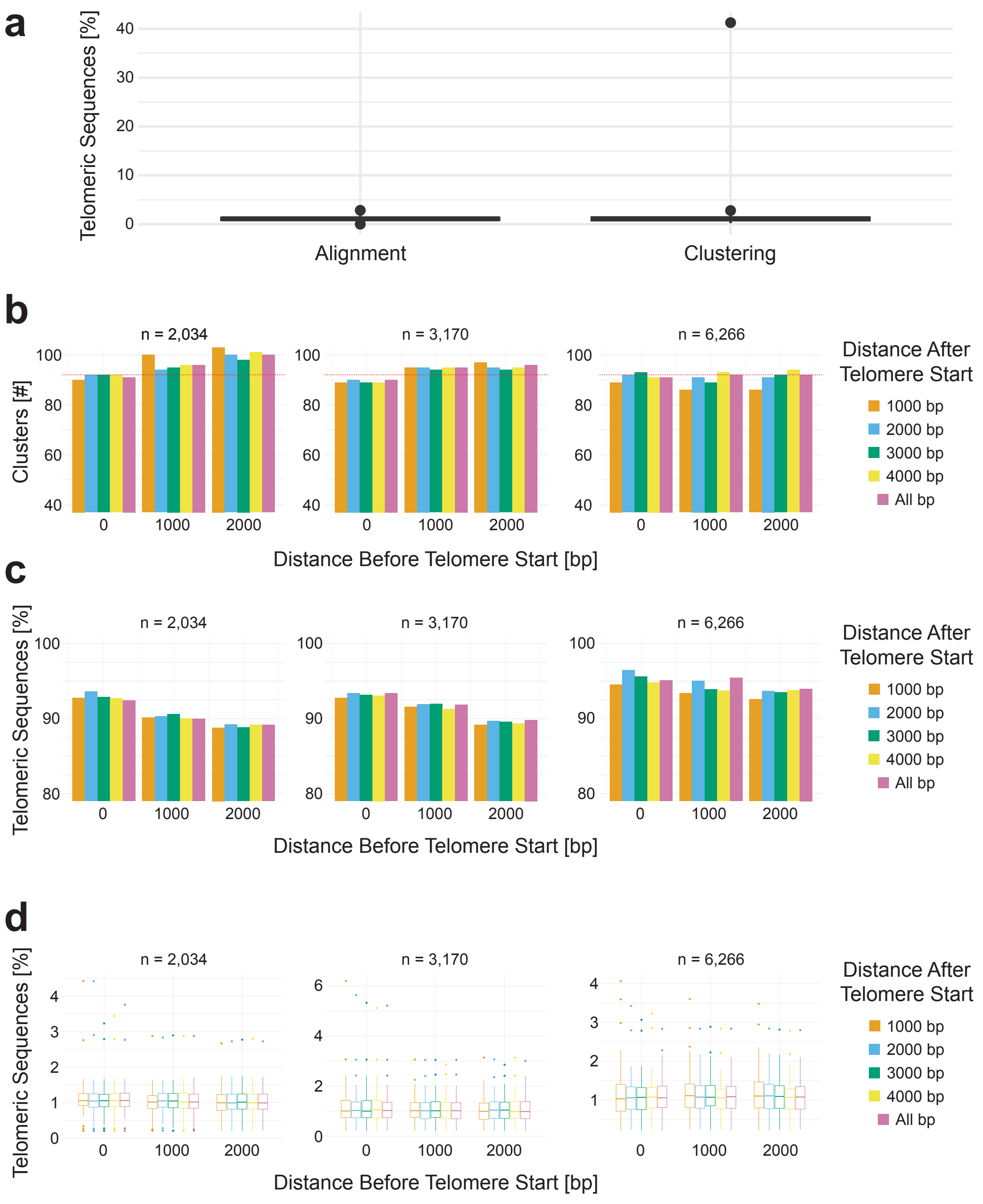


**Deimler, 2025; Supplemental Figure 6**

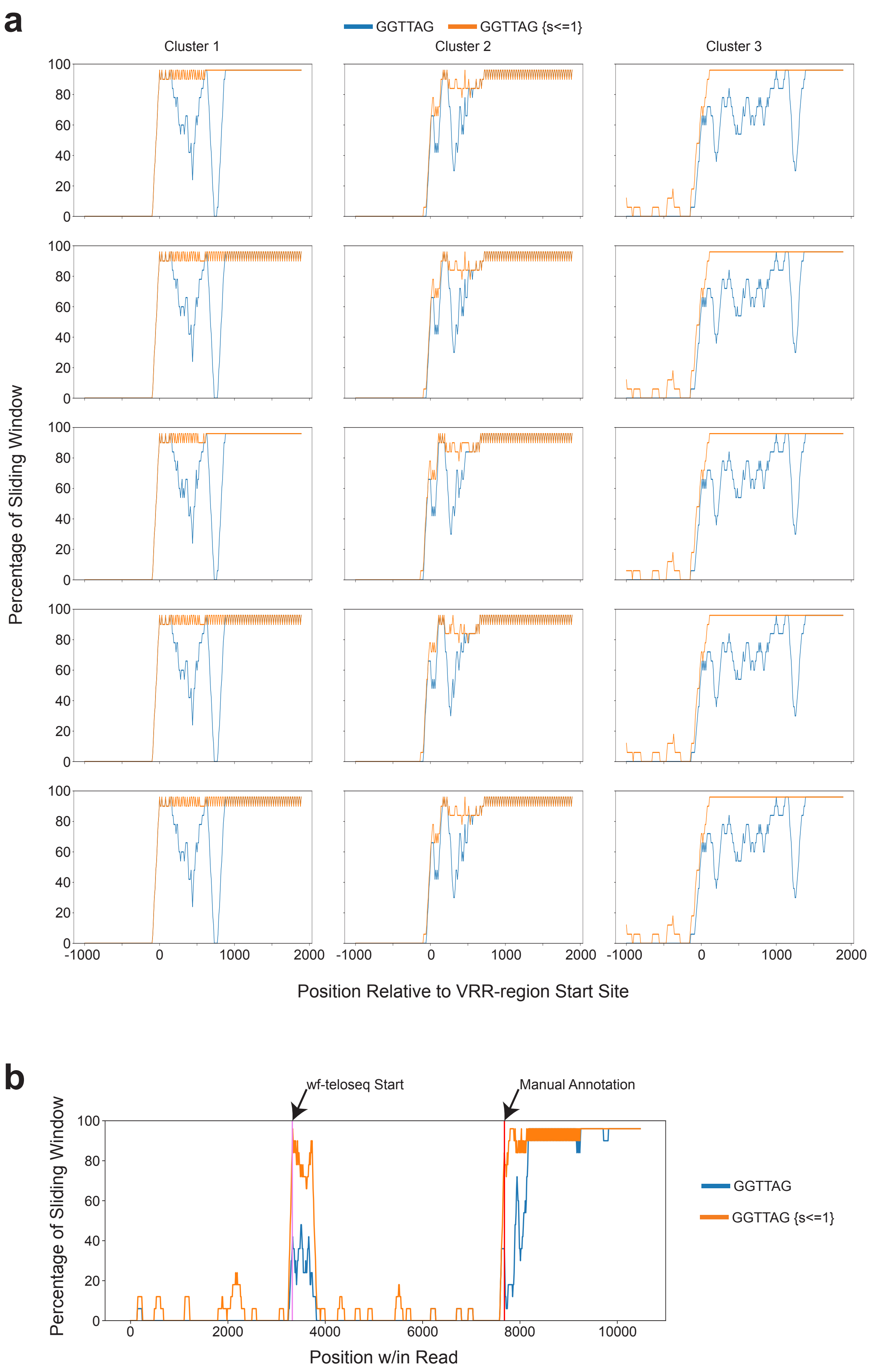


**Deimler, 2025; Supplemental Figure 7**
